# A General Strategy to Endow Dyes with Genic Fluorescence for Wash-Free Imaging of Proteins in Living Cells

**DOI:** 10.1101/2024.12.26.630335

**Authors:** Xuelian Zhou, Lu Miao, Ning Xu, Yonghui Chen, Jinjing Shi, Jinhua Zhou, Guangying Wang, Qinglong Qiao, Zhaochao Xu

## Abstract

Fluorescent dyes are the main fluorophore for protein labeling and fluorescent imaging in living cells. However, due to the inevitable nonspecific binding of the dye to non-targets within the cell, background signals are generated, which severely affect the imaging quality. Here, we endow the dye with genic fluorescence by introducing a fluorescent proteins FRET pair, such that only the dye bound to the target protein emits fluorescence (λ_ex_ = FPs, λ_em_ = Dyes), while nonspecifically bound dye remains non-fluorescent. This significantly improves the signal-to-noise ratio (SNR) in wash-free imaging. This strategy is achieved by fusing the Halo-tag to fluorescent proteins (sfGFP, mCherry) that can generate FRET with Halo dyes **O-Rho** and **Si-Rho**. The FPs serve as the donor, and the FRET mechanism imparts genic fluorescence property to **O-Rho** and **Si-Rho**. Since the excitation wavelength of the donor (λ_ex_ = FPs) cannot excite the unspecifically dye fluorescence, background fluorescence interference is reduced. We further improved the SNR by increasing the FRET efficiency between the FPs and dyes. The system was used for four-color super-resolution imaging of target proteins and dynamic SIM tracking of mitochondria. In addition, the genic FRET fluorophore constructed with the ultra-stable mStayGold showed significant photobleaching resistance and can be used for long-term dynamic super-resolution imaging. Theoretically, by selecting appropriate FRET pairs, this method can improve the SNR of any fluorophore, providing a new labeling strategy for live-cell wash-free imaging.

## 1. Introduction

The combination of appropriate protein fluorescent labeling strategies and advanced fluorescence microscopy is a powerful tool for high-resolution visualization of protein distribution and function in live cells ^1,2^ Small molecule dyes have become the main fluorophores in protein labeling and super-resolution imaging due to their high brightness, photostability and easy chemical modification.^3–5^ Self-labeling protein tags (such as Halo-tag/SNAP-tag) enable the specific labeling of fluorescent dyes to target proteins.^6–8^ However, as exogenous species, it is difficult to control the distribution and quantity of fluorescent dyes in cells, and unspecific binding will inevitably occur to generate background fluorescence.^9,10^ Even with prolonged and multiple washings, these background signals cannot be completely eliminated. This severely affects the quantitative tracking of target proteins, especially low-abundance proteins, and the issue becomes even more prominent as the resolution reaches the single-molecule level.^11–13^

To address this challenge, researchers tried to develop the fluorogenic dyes with genetic encoding and in situ fluorescence generation properties similar as fluorescent proteins. For instance, various protein tag-based "fluorogenic" dyes have been designed, which activate fluorescence upon recognition by the tag protein.^14,15^ However, the binding of these dyes to non-specific targets in living cells can also turn on the fluorescence of the dye (Scheme 1a,b). As a result, although these dyes exhibit high fluorescence turn-on ratios in vitro, they show a lower signal-to-noise ratio (SNR) in live-cell imaging.^16–18^ Taking the widely used rhodamine-based probes as an example, the fluorescence off-on states of these probes is achieved by modulating the balance between the non-fluorescent spirolactones (L) and the fluorescent zwitterions (Z) within the rhodamine structure. When binding to the protein tag, the microenvironment of rhodamine changes which shifting the spirocyclization equilibrium from L to Z and turning on fluorescence.^19,20^ However, the L-Z conversion of rhodamine is not only induced by binding with the protein tag but also by intracellular environmental factors, such as pH,^21^ polarity,^22^ metal ions,^23^ and hydrophobic cavities of proteins,^24^ which induce background signals of rhodamine dyes (such as non-specific binding to mitochondria and lysosome) (Scheme 1b).^25,26^ Recently, Wu *et al.* developed typical fluorogenic probes that, through bioorthogonal reactions, enables the in situ generation of fluorophores with large Stokes shifts within cells.^27^ However, how to enable any dye to form new fluorescent properties in situ during protein recognition remains a challenge.

In this work, we developed a general strategy to endow exogenous fluorescent dyes with genic fluorescent properties, which improves the SNR of dyes in no-wash live-cell imaging. The strategy is achieved by fusing protein tag with a fluorescent protein (FP)^28^ which can produce förster resonance energy transfer (FRET)^29^ with the tag dyes (Scheme 1c). In this system, the fluorescent protein serves as an energy donor, and endows fluorescent dyes with genic fluorescent properties (λ_ex_ = FPs, λ_em_ = Dyes) through FRET. Since the excitation wavelength of the donor (λ_ex_ = FPs) cannot excite the unspecific dye fluorescence, background fluorescence interference is reduced, thereby enhancing the SNR in live-cell protein imaging without washing.

To be specific, we fused Halo-tag with green fluorescent protein (GFP) and red fluorescent protein (RFP) respectively, where the fluorescent proteins as donor and Halo dyes **O-Rho** and **Si-Rho** as receptor (Scheme 1 right), constructing two types of genic fluorophores GLH-O and CLH-Si. This approach improved the fluorescence SNR of the weak-fluorogenic **O-Rho** and fluorogenic **Si-Rho** dyes under lower protein expression level. To further improve the SNR, we shortened the linker between mCherry and Halo-tag to enhance FRET efficiency. The obtained genic fluorophore CH-Si enabled no-wash four-color super-resolution imaging of target proteins and the tracking of dynamic interactions between mitochondria and endoplasmic reticulum/lysosomes. Finally, we applied the photostable green fluorescent protein mStayGold to develop a genic fluorophore SLH-O with excellent photostability. This strategy can improve the SNR of any fluorescent dyes and provide a new technical tool for real-time, quantitative, and long-term super-resolution dynamic tracking of target proteins.

**Scheme 1.**
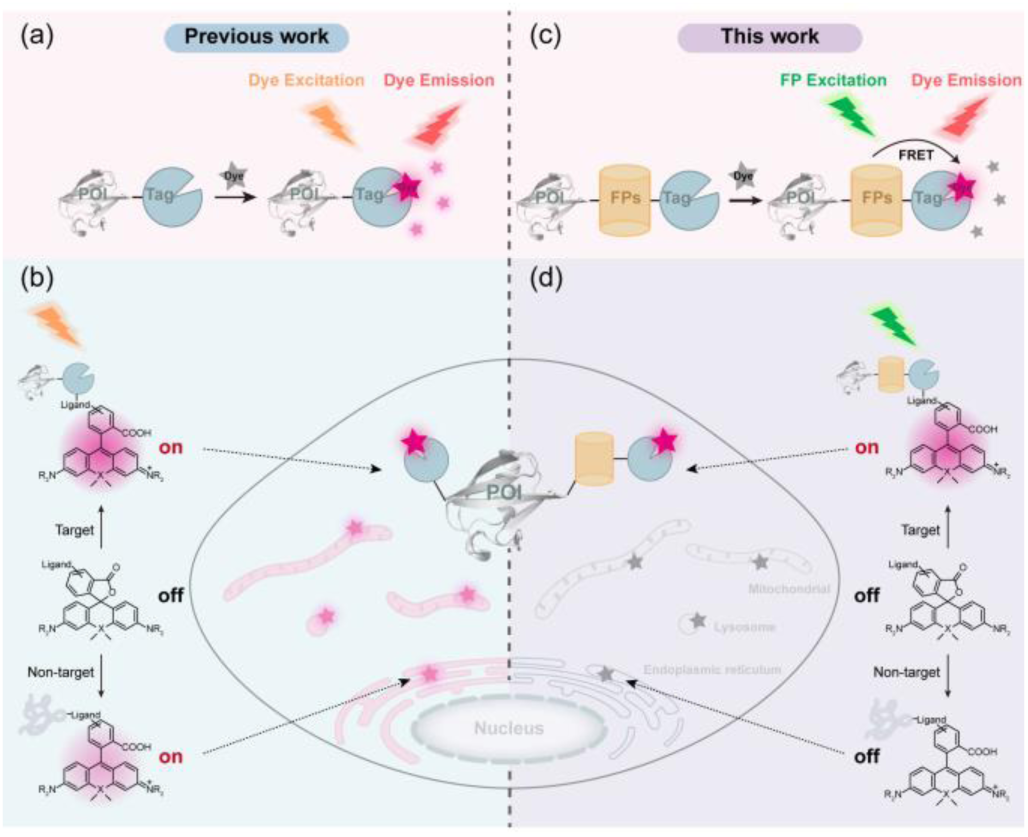
Comparison of fluorogenic probe design strategies based on protein tags and FRET. (a) traditional fluorogenic probes of self-labeling protein tags. (b) The environment-dependent switch between the spirolactones (L) and zwitterions (Z) in Rhodamine is prone to nonspecific background signals. (c) The genetically coded fluorophores were constructed by fusing the Halo-tag to fluorescent proteins. (d) it avoids direct excitation of non-specifically dye fluorescence and effectively reduce background fluorescence.

## 2. Results and discussion

### 2.1 Evaluation of Halo-tag-based dyes

Rhodamine (Rho), as a traditional fluorescent dye, has been widely used for the labeling and imaging of proteins in live cells and tissues.^29^ Following the previously described synthetic method, we respectively conjugated oxyrhodamine and silicon rhodamine with the chloroalkyl group ligand to obtain two kinds of Halo-tag probes, **O-Rho and Si-Rho**.^30,31^ Spectroscopic characterization in vitro showed different changes in absorption and fluorescence turn-on for both probes, with and without the presence of the Halo-tag. **O-Rho** mainly exists in the form of fluorescent zwitterions in aqueous solution. After binding to Halo-tag, the fluorescence emission intensity at 569 nm increased by only 1.6-fold (Figure 1a and Figure S1a). In contrast, **Si-Rho** mainly exists in the form of non-fluorescent spironolactone due to aggregation in aqueous solution, which exhibited a 6.2-fold increase in fluorescence emission at 662 nm upon binding to Halo-tag (Figure 1c and Figure S1b). This result was basically consistent with previous reports.^17^

**Figure 1.**
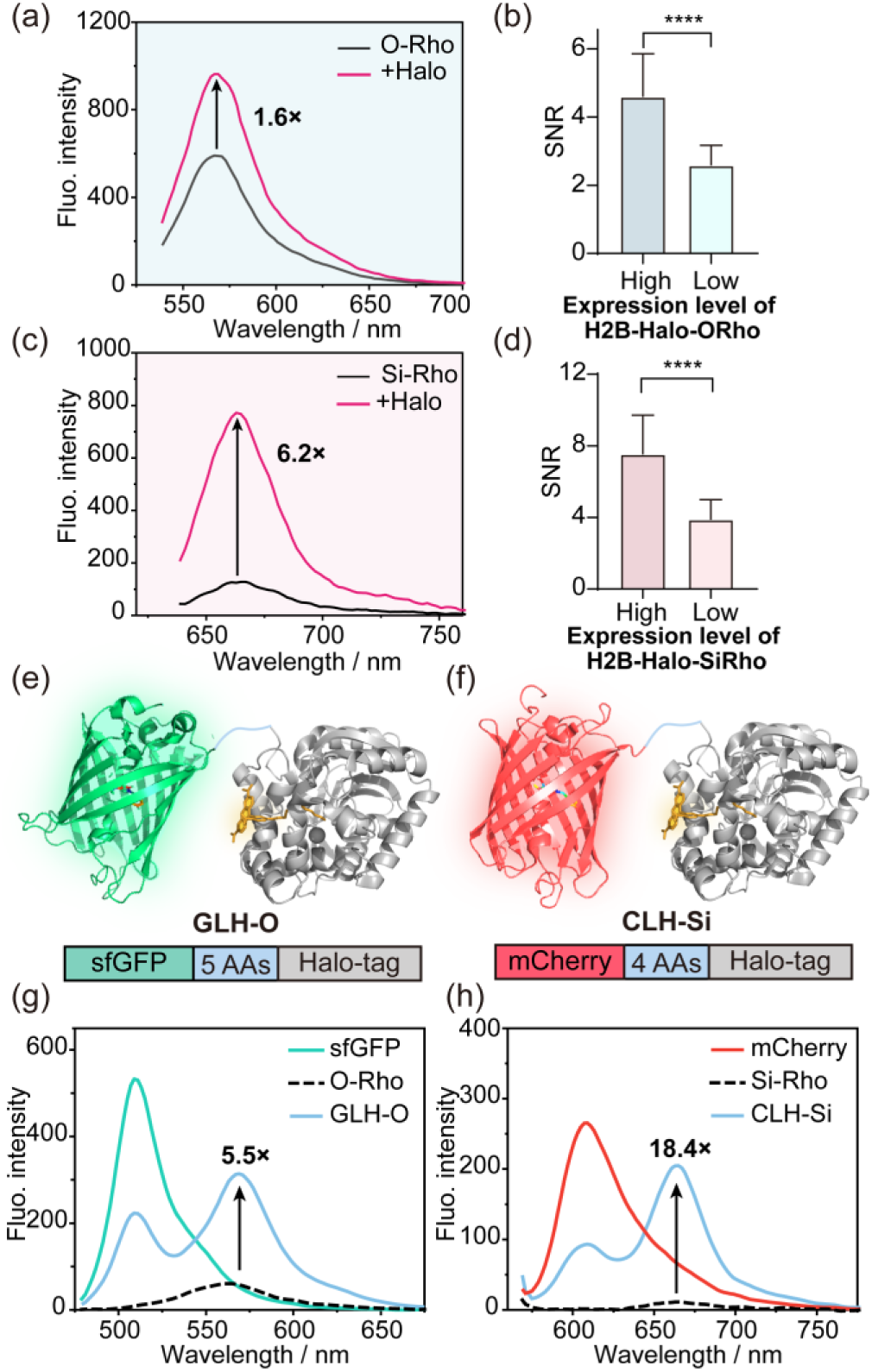
Characterization of traditional fluorogenic probes and genic FRET probes. (a, c) Emission spectra of **O-Rho** (a) and **Si-Rho** (c) in the presence and absence of excess Halo-tag. **O-Rho:** λex= 530 nm, **Si-Rho:** λex: 630 nm. (b, d) At the same dye labeling concentration (**O-Rho**: 0.1 μM; **Si-Rho**: 0.3 μM), the difference of SNR with varying H2B-Halo protein expression levels in HeLa cells. Data are means ± S.D.; n = 40-50 cells; Each group represents 3 independent experiments separately; Unpaired two-tailed Student’s t-test, ****p≤0.0001. (e, f) Schematic representation of the GLH-O and CLH-Si structure of Halo-tag (PDB: 6y7a) fused at the C-terminal of sfGFP (PDB: 2B3P) and mCherry (PDB: 2H5Q). (g) Fluorescence emission spectra of **O-Rho** with and without the addition of GLH (λex: 470 nm).GLH-O. (h) Fluorescence emission spectra of **Si-Rho** with and without the addition of CLH (λex: 561 nm).

Subsequently, we used **O-Rho** and **Si-Rho** to directly stain living cells (Figure S2a and S3a). The fluorescence of mitochondria and lysosomes was significantly enhanced in both **Si-Rho and O-Rho** treated cells, which indicating that non-specific binding could induce a shift in the spirocyclization equilibrium from L to Z and turn on fluorescence (Scheme 1b). As the dye concentration increased, the non-specific background signal also grew stronger (Figure S2b, and S3b). We then incubated these probes with Hela cells over-expressing H2B-Halo protein (Figure S2c, and S3c). As expected, with the increase of dye concentration, the SNR (fluorescence intensity ratio of nuclear to cytoplasmic) decreased significantly (Figure S2d, and S3d). Importantly, at the same dye concentration, cells with relatively lower expression levels of H2B-Halo protein exhibited a lower SNR (Figure 1b, d). However, due to the fluorescence turn-on characteristics of **Si-Rho**, its SNR is higher than that of cells stained with **O-Rho** dye. The **O-Rho** probe showed a certain SNR (higher than that in vitro) because the over-expression of H2B-Halo in the cell nucleus caused a large amount of dye to be concentrated in nucleus. Overall, the turn-on of non-specific fluorescence in fluorogenic fluorescent dyes significantly affects the SNR of live cell imaging.

### 2.2 Design and construction of genic fluorophores

**O-Rho** and **Si-Rho** are two types of Halo-tag probes with different fluorescence emission characteristics, and exhibiting weak-fluorogenic and fluorogenic properties, respectively (Figure 1a, c). We constructed genic fluorophores based on **O-Rho** and **Si-Rho** to demonstrate the universality of this strategy. In order to achieve FRET, we considered factors such as the good overlap of the absorption and emission spectra between the donor and acceptor, the high quantum yield of the donor, and the excellent extinction coefficient of the acceptor (Figure S4).^28^ We finally selected sfGFP and mCherry as donors of **O-Rho** and **Si-Rho** (Figure 1e, f), respectively Halo-tag. Both fusion proteins were expressed and purified in vitro, and SDS-PAGE confirmed that the purified proteins had the correct molecular weight and could be covalently labeled with Halo dyes (Figure S5-6). We obtained genic fluorophores sfGFP-linker-Halo-ORho (hereinafter referred to as GLH-O) and mCherry-linker-Halo-SiRho (hereinafter referred to as CLH-Si).

In vitro spectral characterization of GLH-O showed that the fluorescence of **O-Rho** (λem = 565 nm) was emitted under the excitation wavelength of the donor sfGFP (λ_ex_ = 470 nm). This was accompanied by a decrease in donor intensity and an increase in the acceptor fluorescence intensity, confirming the occurrence of FRET (E_FRET_ =58.1%, Figure 1g). Similarly, the spectroscopic analysis of CLH-Si demonstrated significant FRET with the efficiency of 64.9% (Figure 1h). In contrast, free **O-Rho** and **Si-Rho** dyes in solution exhibited relatively low fluorescence signals at the same excitation wavelengths, resulting in 5.5-fold and 18.4-fold increase in fluorescence at the acceptor emission wavelengths of GLH-O and CLH-Si, respectively. These enhancements significantly outperformed the **O-Rho** (1.6-fold) and **Si-Rho** (6.2-fold) probes (Figure 1).

### 2.3 Wash-free live-cell imaging of genic fluorophores

First, we tested the FRET effect of GLH-O and CLH-Si probes in living cells. We transiently expressed the fusion proteins of H2B-GLH and H2B-CLH (H2B: human histone) in HeLa cells, and performed fluorescence imaging before and after the addition of the dye. Under 488 nm excitation of the donor, GLH had almost no obvious fluorescence in the FRET emission channel (580-653 nm) before labeling with **O-Rho** dye (Figure S7a). After the addition of the **O-Rho**, the fluorescence in the donor channel (500-550 nm) was significantly weakened, and the signal in the FRET channel was enhanced, which proving the effective occurrence of FRET in GLH-O. Similarly, we also observed FRET signal in HeLa cells expressing H2B-CLH-Si (excitation 561 nm, collection 663-703 nm, Figure S7b) corresponding to the results in vitro.

Genetically encoded FPs act as the donors of Rho dye, causing a blue-shift in the excitation wavelength of Rho dye labeled to the target. Ideally, the fluorescence of the Rho dye not labeled to the target would not be excited, thus reducing the imaging background (Figure 2a, b). To evaluate the no-wash imaging effect of genic fluorophores, we performed imaging at low protein expression levels. We transiently transfected HeLa cells with H2B-GLH and H2B-CLH, and incubated them with 2 μM **O-Rho** and **Si-Rho** dyes after 12 hours of expression for no-wash imaging. The imaging results of single cells showed that the background fluorescence with short wavelength excitation (GLH**-**O: λ_ex_ = 488 nm; CLH-Si: λ_ex_ = 561 nm) was lower than that of the conventional wavelength excitation of the dye (GLH-O: λ_ex_ = 561 nm; CLH-Si: λ_ex_ = 640 nm). The background was reduced by 3.6 and 3.1 folds, respectively, and thereby improving the SNR of the dye (Figure 2a, b). Due to some crosstalk between the FRET pairs, short-wavelength may also excite a certain amount of non-specific fluorescence, so the background fluorescence has not been eliminated.

**Figure 2.**
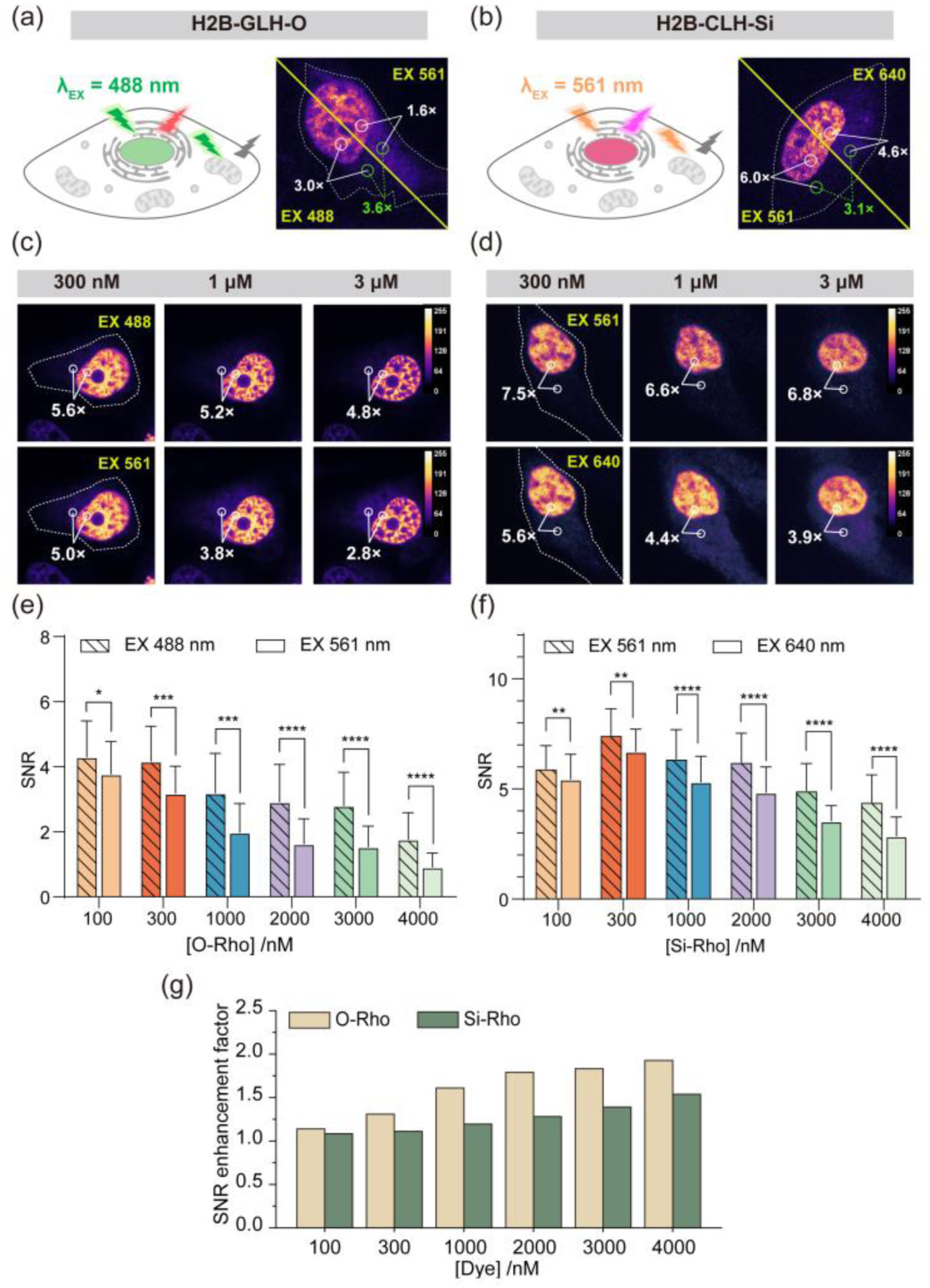
Wash-free live-cell imaging of genic fluorophores. (a, b) schematic diagram and the wash-free confocal imaging of genic fluorophores GLH-O and CLH-Si [**O-Rho**]: 2 μM; [**Si-Rho**]: 2 μM. (c, d) Comparison of no-wash imaging in the FRET channel and the acceptor channel for H2B-GLH-O (c) or H2B-CLH-Si (d) protein labeled with varying concentrations of Rho. Calibration bar shows the normalized fluorescence intensity. (e, f) Statistical analysis of the SNR in the FRET channel and Rho channel for HeLa cells expressing GLH-O (e) or CLH-Si (f) under different dye concentration treatments. Data are means ± S.D.; n = 40-50 cells; Each group represents 3 independent experiments separately; Unpaired two-tailed Student’s t-test, *p ≤ 0.05, **p ≤ 0.01, ***p ≤ 0.001, ****p ≤ 0.0001. (g) The enhancement factor of the mean SNR in the FRET channel compared to the dye channel for GLH**-**O and CLH-Si probes, labeled with different concentrations of Rho dye.

We further quantified and compared the differences of SNR in the same cell after incubation with various concentrations of **O-Rho** and **Si-Rho** dyes (0.1 μM-4 μM). The results demonstrated that the SNR in FRET channel was higher than that in the dye channel under all **O-Rho**/**Si-Rho** dye concentrations (Figure 2c-f). Additionally, as the dye concentration increased, the SNR improved more, and the SNR of GLH-O showed a greater increase compared to CLH-Si (Figure 2g). Unfortunately, due to the influence of FRET efficiency and crosstalk between the FRET pairs, the overall SNR enhancement was limited to 1.5–2 fold. In theory, if a perfect FRET pair without crosstalk is used, non-specific background signals can be completely eliminated to obtain ideal genic fluorescence.

### 2.4 The improvement of FRET efficiency further enhanced the SNR of genic fluorophores

The CLH-Si probe showed higher SNR, and the near-infrared emission, making it potentially useful for biological imaging in complex systems. To further enhance its SNR, we considered improving the FRET efficiency of CLH-Si. Although background signal based on fluorescence crosstalk of mCherry-SiRho FRET pair is unavoidable, increasing FRET efficiency can increase the brightness of the genic fluorophore, thereby improving its SNR. We truncated the linker amino acids between mCherry and Halo-tag fusion protein to obtain mCherry-Halo (hereinafter referred to as CH, Figure 3a). The fusion protein was expressed, purified, and then labeled with **Si-Rho** to generate CH-Si (Figure S6). The spectral characterization showed significantly enhanced energy transfer compared with CLH-Si, and the FRET efficiency was increased from 64.9% to 83.7% (Figure 3b). Under 561 nm excitation, CH-Si exhibited a 25-fold increase in fluorescence at the dye emission wavelength (662 nm) compared to free **Si-Rho** in solution.

**Figure 3.**
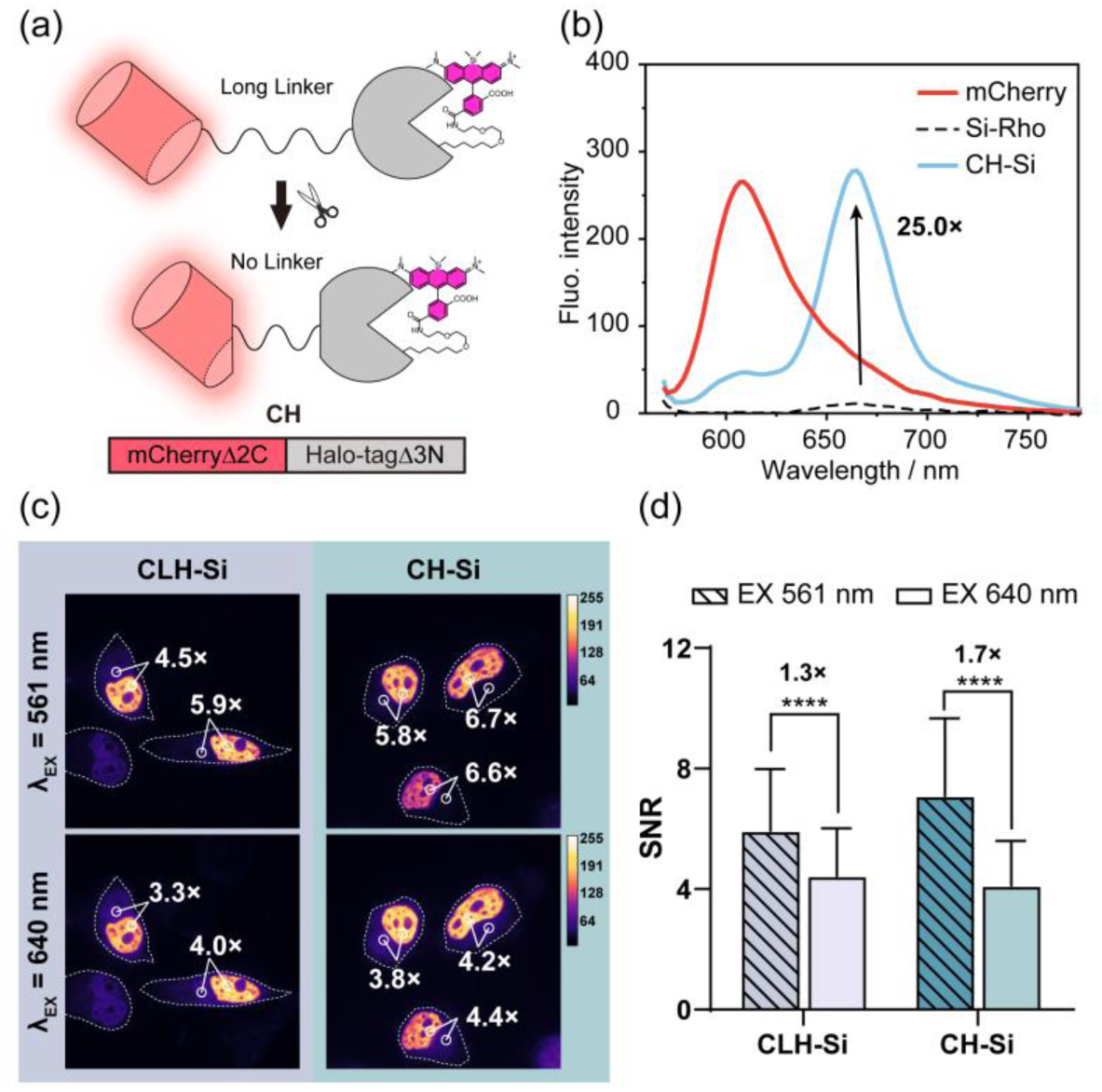
Development of genic fluorophores with higher SNR. (a) Protein engineering schematic of CH fluorogenic probe. (b) Fluorescence emission spectra of CH protein with (blue line) and without (red line) the addition of **Si-Rho** (λex: 561 nm). The numbers indicate the ratio between the fluorescence of CH-Si and free **Si-Rho** (black dot line) at 662 nm (λex: 561 nm). (c) Comparison of no-wash imaging in the FRET channel and the **Si-Rho** channel for H2B-CLH-Si or H2B-CH-Si. [**Si-Rho**]: 2 μM. Calibration bar shows the normalized fluorescence intensity. (d) Statistical analysis of the SNR in the FRET channel and Rho channel from the images of (c). Data are means ± S.D.; n = 40-50 cells; Each group represents 3 independent experiments separately; Unpaired two-tailed Student’s t-test, ****p ≤ 0.0001.

Furthermore, we transiently transfected Hela cells with CH-H2B. After 12 hours of expression, the cells were labeled with 2 uM **Si-Rho** and used for no-wash imaging. The SNR of fluorescence imaging in the same cell at 561 nm excitation was improved compared to that at 640nm excitation (Figure 3c). The results compared with CLH-Si showed that the SNR of genic fluorescence increased from 4.87±1.68 to 5.92±1.32 after the modification, while the SNR of dye channel remained basically unchanged (CLH-Si: 3.59±0.792; CH-Si: 3.90±0.483). The overall SNR was increased from 1.3-fold to 1.7-fold (Figure 3d). As expected, CH-Si exhibited a higher SNR, confirming the effectiveness of the FRET optimization.

### 2.5 The universality of CH-Si for labeling different proteins in living cells

In previous experiments, CH-Si probe was successfully used labeling H2B in HeLa cells (Figure S8a). Following the plasmid construction method described in the SI, we continued to fuse CH to the N terminus of mitochondrial outer membrane protein TOMM20, endoplasmic reticulum tail-anchored protein Sec61β, and GTPase Rab7 (targeting lysosome). After 12 hours of transient transfection, no-wash fluorescence imaging was performed. The imaging results showed that CH-Si can clearly label different cellular organelles (Figure S8b-d), and the dual protein fusion system will not interfere with the correct localization of these subcellular structural proteins. Furthermore, CH-Si exhibited good SNR in imaging different organelles (Figure S8e). These imaging results demonstrated the broad potential of CH-Si as a versatile tool for biological labeling applications.

### 2.6 Using CH-Si for no-wash four-color SIM imaging

The study of most biological functions requires the simultaneous observation of multiple biomolecules and their interactions.^32^ Therefore, fluorescent probes for multicolor live-cell imaging must have not only sufficient spectral resolution to distinguish various fluorescently labeled structures, but also characteristics suitable for no-wash imaging to reduce the complexity of sample handling and improve imaging efficiency.^33,34^ The blue-shifted excitation of CH-Si provides it with an extended Stokes shift characteristic (75 nm), enabling it to be used in conjunction with fluorophores of conventional channels for multichannel imaging. Here, we first examined the feasibility of dual-labeling using CH-Si in conjunction with traditional RFPs for confocal imaging. This requires that CH-Si has almost no residual fluorescence in the donor emission channel. We labeled H2B protein in HeLa cells with CH-Si and mCherry proteins, respectively, and quantified the fluorescent signals from donor channel (λ_ex_ = 561 nm, collection 580-653 nm) and FRET channel (λ_ex_ = 561 nm, collection 663-703 nm). By calculating the fluorescence ratio of the two channels, we determined the residual fluorescence level of CH-Si in the donor channel (Figure S9a). The results showed that the fluorescence ratio of CH-Si was significantly reduced compared with mCherry. This indicates that CH-Si has lower fluorescence in the donor channel and stronger FRET emission, which can be effectively distinguished from RFPs. Figure S9b showed dual-color imaging in HeLa cells using mCherry and CH-Si, where lysosomes (green channel) and mitochondria (red channel) subcellular organelles can be distinguished by different colors without obvious fluorescence interference.

Structured illumination microscopy (SIM) is a powerful imaging technique that can provide high-resolution images of biological structures in living cells.^35^ Similar to traditional microscopy, the application effect of SIM in living-cell is largely limited by the types and number of available fluorescent probes, which directly affecting its performance and application in multi-color imaging.^36^ First, we compared the SNR of CH-Si (λ_ex_ = 561 nm, λ_em_ = 700 ± 75/2 nm) and Halo-Si (λ_ex_ = 640 nm, λ_em_ = 663–738 nm), showing that CH-Si also has a better SNR in SIM super-resolution multicolor imaging (Figure 4a-b). Then we combined CH-Si with several traditional fluorophores for four-color imaging, where CH-Si was labeled on TOMM20, Hoechst 33342 was stained on the nucleus, EGFP was fused to Sec61β, and mCherry was labeled on Rab7 (Figure 4c).

**Figure 4.**
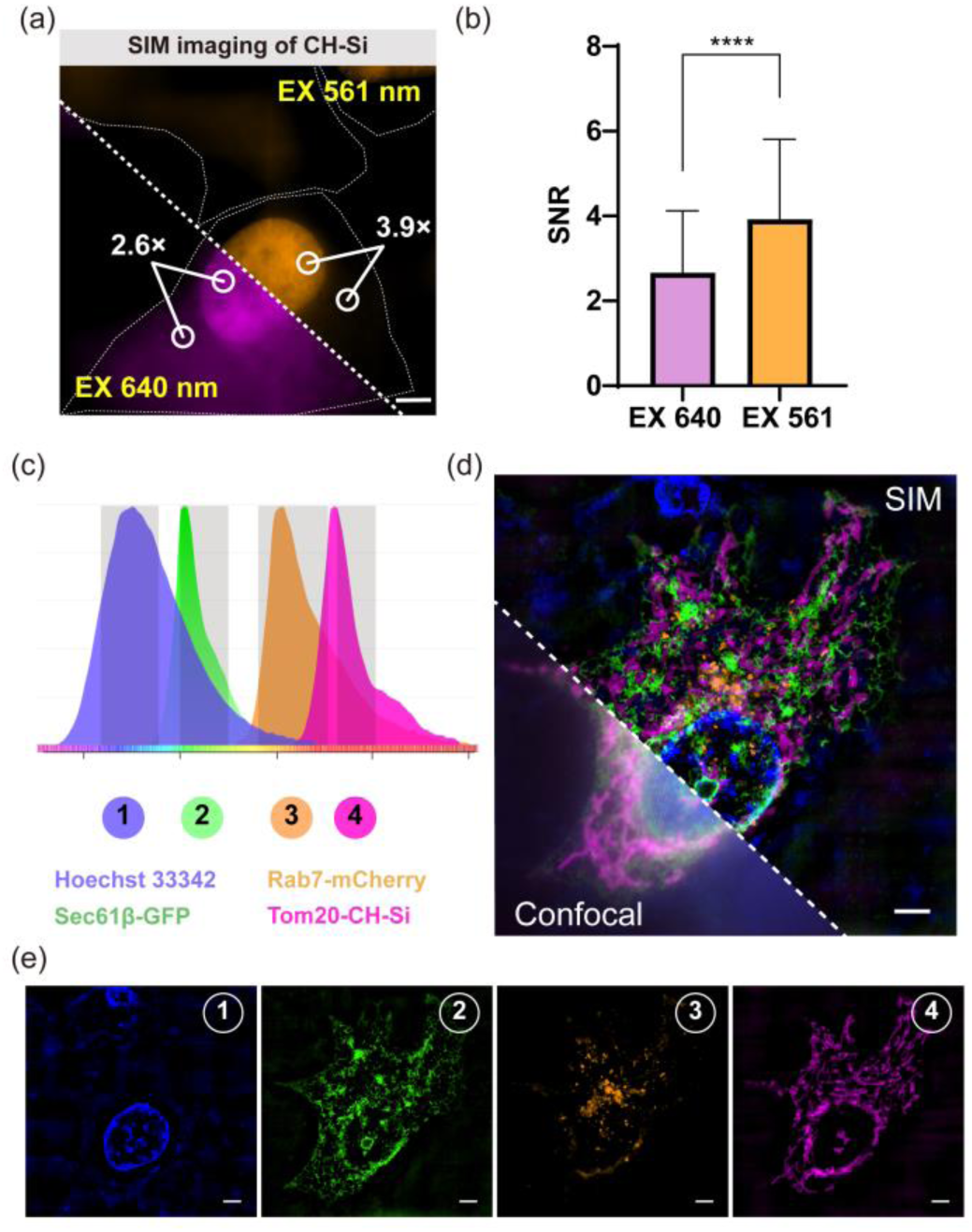
CH-Si for wash-free live-cell SIM imaging. (a) Comparison of no-wash SIM imaging in the FRET channel and the **Si-Rho** channel for H2B-CH-Si. [**Si-Rho**]: 2 μM. Scale bar: 5 μm. (b) Statistical analysis of the SNR in the FRET channel and Rho channel from the images of (a). Data are means ± S.D.; n = 40-50 cells; Each group represents 3 independent experiments separately; Unpaired two-tailed Student’s t-test, ****p ≤ 0.0001. (c) Emission spectra of the different fluorophores used for four-color imaging. The grey area shows the collection wavelength range from the different channels, which marked with different color numbers. (d) Four-color confocal and SIM images of nucleus stained with Hoechst 33342 (blue ①), Sec61β labeled with EGFP (green ②), Rab7 labeled with mCherry (orange ③), and TOMM20 labeled with CH-Si (pink ④). (e) The bottom panels show the separated channel images corresponding to the overlay in (b). Scale bar: 5 μm.

By using these four different fluorophores, we successfully achieved SIM imaging of four substructures (Figure 4d-e): the nucleus (blue ①), endoplasmic reticulum (green ②), lysosome (orange ③), and mitochondria (pink ④). These results suggested that the combination of CH-Si with other fluorescent probes provided a powerful tool for monitoring the dynamics of multiple organelles.

### 2.7 SIM dynamic imaging of mitochondria with CH-Si

The dynamic behavior of various biomolecules and organelles in cells is often extremely complex and rapidly changing. Precise positioning and high-resolution imaging are essential for accurately capturing these dynamic processes.^37^ The position and movement of mitochondria in living-cells play a key role in cellular energy metabolism, signal transduction, and apoptosis.^38,39^ Mitochondria carry out their functions through a close network of interactions with other organelles such as the endoplasmic reticulum (ER) and lysosomes.^40,41^ In this study, CH-Si based SIM imaging was used to track the dynamics interactions of mitochondria with the ER and lysosomes.

To capture the interaction between mitochondria and ER, we co-expressed TOMM20 protein labeled with CH-Si and EGFP-Sec61β in Hela cells. We performed dual-color SIM imaging to continuously track the dynamics of mitochondria and ER for 20 minutes (Figure 5a-c). The boxes highlight the details of the interaction between mitochondria and ER at different locations. In Figure 5b, we observed a significant morphological change in mitochondrion at 100 s, indicated by the blue arrow, where it elongated and briefly contacted the adjacent ER. Over time (110-160 s), the mitochondrial morphology gradually recovered, shrank and broke away from the contact with ER. The interaction between mitochondria and ER is not limited to physical contact, but also has an important impact on the function and morphology of mitochondria. In Figure 5c, we found that the mitochondria underwent fission process (300-310 s) after a rapid interaction with the ER (270 s). The combined labeling of CH-Si and EGFP can reveal the complex and delicate dynamic relationship between mitochondria and endoplasmic reticulum.

**Figure 5.**
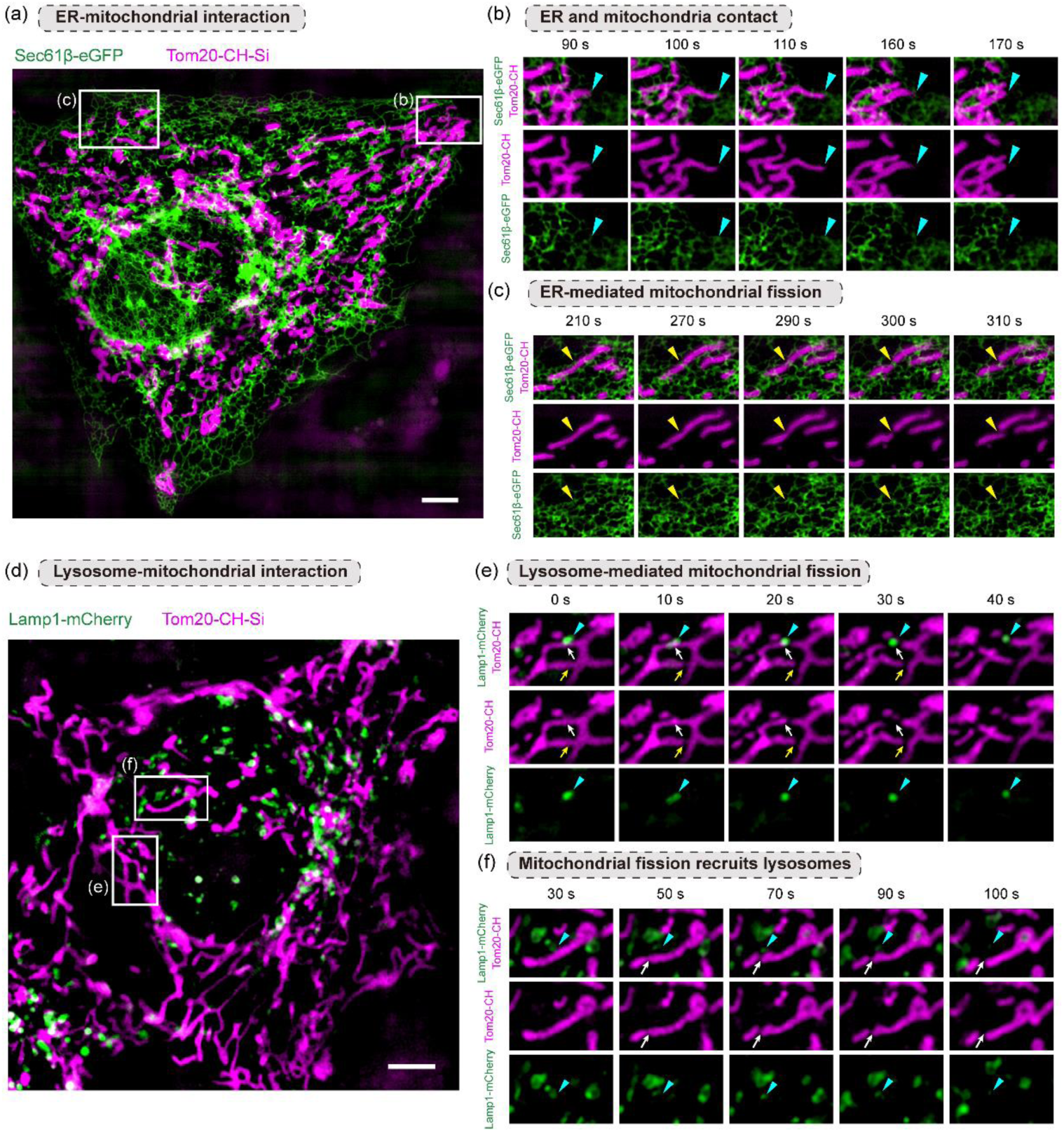
Dual-color SIM imaging of mitochondria dynamic using CH-Si. (a) Dual-color SIM image of HeLa cells co-expressing GFP-Sec61β and TOMM20-CH-Si. The endoplasmic reticulum and mitochondria are represented in green and pink channels, respectively. Scale bar: 5 μm. (b) Magnified images of the boxed region in (a). The blue arrows indicate the contact event between mitochondria and the endoplasmic reticulum. (c) Magnified images of the boxed region in (a). The yellow arrow indicates the location where endoplasmic reticulum-induced mitochondrial fission event occurred. (d) Dual-color SIM image of HeLa cells co-expressing mCherry-Rab7 and TOMM20-CH-Si. The lysosome and mitochondria are represented in green and pink channels, respectively. Scale bar: 5 μm. (e, f) Magnified images of the boxed region in (d). The two groups of images represent the two modes of action of lysosomes in the fission of mitochondria.

To further track the dynamic interaction between mitochondria and lysosomes, we used mCherry to label the lysosome protein Rab7. We successfully demonstrated the high-resolution separation of the two fluorescence signals, ensuring the resolution accuracy of mitochondria and lysosomes (Figure 5d-f, Figure S9b). Figure 5e shows the dynamic process of lysosomes participating in mitochondrial fission. The lysosome indicated by the blue arrow induced the fission of the mitochondria (30 s) after contacting the mitochondria (10 s). This indicates that the lysosomes play a significant role in the process of mitochondrial fission.^42^ However, in the adjacent key region indicated by the yellow arrows, mitochondria did not exhibit significant interaction with lysosomes throughout the observation period, suggesting that lysosomes are not always necessary for the fission of certain mitochondria. This implies the existence of different mitochondrial fission patterns. In addition, we observed another interaction pattern between mitochondria and lysosomes. In Figure 5f, after the mitochondria indicated by the white arrow completed the fission (50-70 s), the lysosomes indicated by the blue arrow quickly moved to the fission site at 70 s and remained in the area for a period of time, possibly carrying out important molecular transport or other biological processes.^42^ Subsequently, the lysosomes moved out of the the fission site at 90 s. This phenomenon may indicate that lysosomes perform certain functional tasks on post-fission mitochondrial, such as membrane repair, material exchange or signal transduction.^43^

### 2.8 Genic fluorophores with high photostability

To achieve long-term super-resolution imaging, it is crucial to develop genic fluorophores with excellent photostability. However, traditional fluorescent proteins exhibit relatively low photostability, and their combination with small molecule dyes will reduce the photostability of the dyes. To address this, we selected the recently developed highly photostable monomeric green fluorescent protein mStayGold^44^, and fused it with the Halo-tag tag. In conjunction with the **O-Rho** dye (Figure 6a), we constructed a new genic fluorophore mStayGold-Linker-Halotag-**ORho** (hereinafter referred to as SLH-O). We tested the photostability of SLH-O through long-term SIM super-resolution imaging. The results demonstrated that SLH-O exhibited significantly enhanced photostability compared to GLH-O (Figure 6b-c), and SLH-O also showed excellent resistance to photobleaching under strong laser irradiation (488 nm, 0.16 kW/cm^2^). This finding proves that the combination of mStayGold with **O-Rho** dye can significantly enhance the photostability of genic fluorophores, providing a new tool for long-term dynamic tracking and no-wash labeling in complex systems.

**Figure 6.**
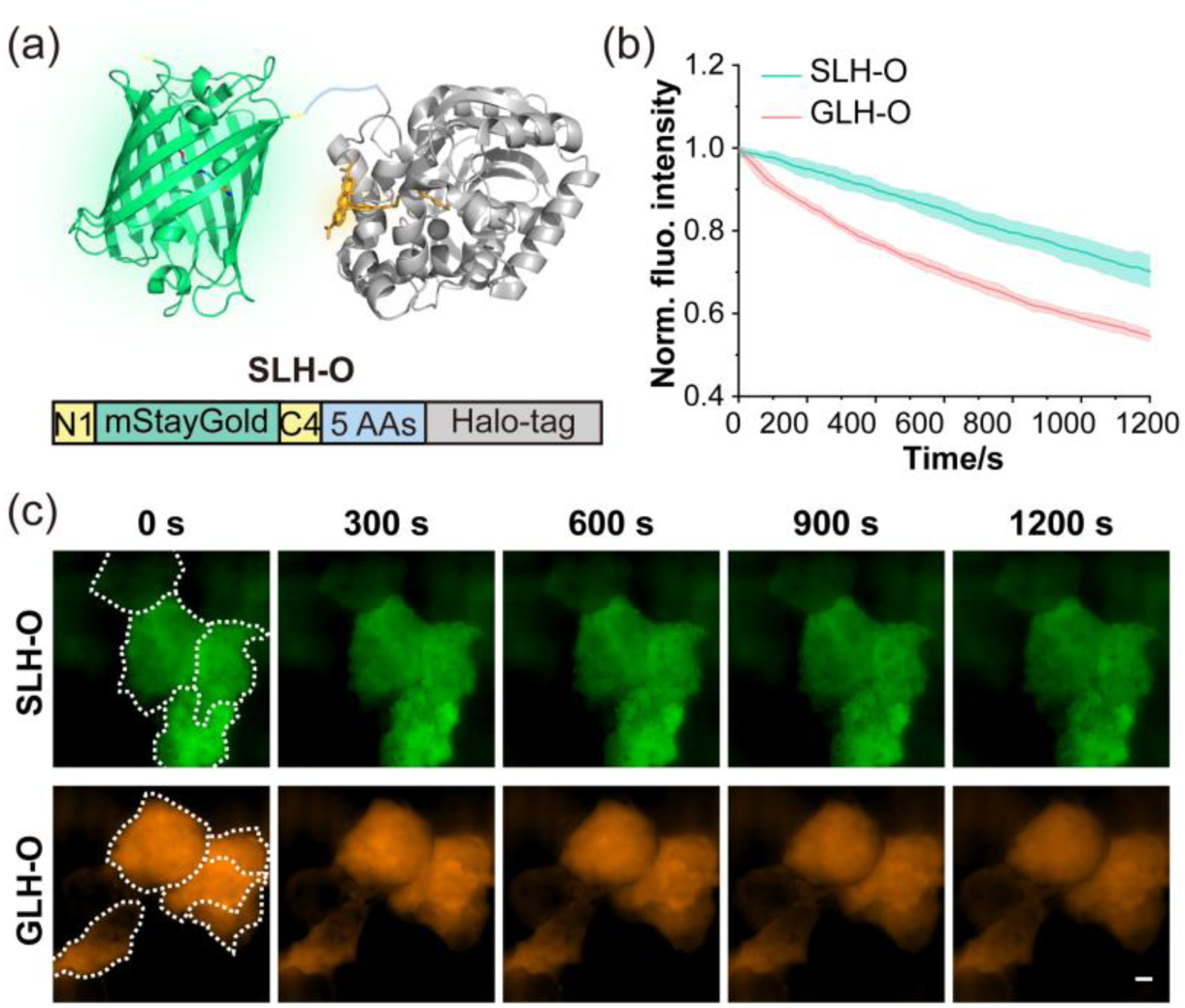
Construction and characterization of photostable genic fluorophores. (a) Schematic representation of the structure of Halo-Tag (PDB: 6y7a) fused at the C-terminal of mStaygold (PDB: 8BXT). (b) Photobleaching dynamics comparison of SLH-O and GLH-O with SIM super-resolution imaging. Each curve represents the average of three independent experiments. Laser export power: 488 nm, 0.16 kW/cm^2^. (c) SIM photobleaching images over time of SLH-O and GLH-O expressed in the cytoplasm.

## 3. Conclusions

In this paper, we successfully developed a universal strategy to convert exogenous fluorescent dyes into probes with genic fluorescence, which significantly improved the SNR of fluorescent dyes in no-wash super-resolution imaging of living cells. We validated the effectiveness of this strategy by applying genic fluorophores (GLH**-**O, CLH-Si) in HeLa cells, and found that the SNR in FRET channel was higher than that in the dye channel under all **O-Rho**/**Si-Rho** dye concentrations (0.1 μM-4 μM). Additionally, the improved FRET efficiency (83.7%) in the CH-Si system effectively reduces background fluorescence, providing clearer and more accurate imaging of subcellular structures, and improving the imaging SNR from 1.3-fold to 1.7-fold. In addition, we successfully combined CH-Si with other fluorophores to achieve four-color SIM imaging and demonstrated the dynamic interactions between mitochondria and the endoplasmic reticulum/lysosomes through two-color no-wash imaging. Furthermore, by introducing the SLH construct, we achieved long-term super-resolution imaging stability. However, challenges remain, as the overall SNR enhancement of CH-Si and GLH-O is limited to 1.5-2 folds due to fluorescence bleed-through. Despite these, the SLH-O and CH-Si probes still show great potential in improving the capabilities of live-cell fluorescence imaging, this strategy enables fluorophores to maintain high signal stability and imaging quality during protein labeling and tracking, providing a reliable tool for long-term super-resolution tracking of complex cell dynamic processes.

## CRediT authorship contribution statement

**Xuelian Zhou:** expressed the proteins, did the cell imaging, and wrote the paper. **Lu Miao:** designed experiments, analyzed the data, and wrote the paper. **Ning Xu:** synthesized the probe. **Qinglong Qiao:** synthesized the probe and examined optical properties. **Yonghui Chen:** expressed the proteins. **Jinjing Shi:** cultured the cells. **Jinhua zhou:** expressed the proteins. **Zhaochao Xu:** designed experiments, and wrote the paper.

## Acknowledgements

This work is supported by the National Natural Science Foundation of China (22225806, 22078314, 22278394, 22378385) and Dalian Institute of Chemical Physics (DICPI202142, DICPI202436).

