## supplemental Materials, methods, and Figures for "A General Strategy to Endow Dyes with Genic Fluorescence for Wash-Free Imaging of Proteins in Living Cells"

### Materials and methods

#### *Materials and instruments.*

Unless otherwise specifically stated, all reagents were purchased from commercial suppliers (Sigma-Aldrich, Innochem, Aladdin, and Sangong Biotech) and used without further purification. All water used was from a Millipore water purification system with a minimum resistivity of 18.0 MΩ · cm. Except for commercially synthesized plasmids (synthesized by General Bio), other plasmids were ordered from the Addgene and NEB website with the corresponding numbers noted. Primers for cloning were synthesized by General Bio in China. UV-vis absorption spectra were collected on an Agilent Cary 60 UV-Vis Spectrophotometer and blank corrected. The blank correction was performed using the respective pure solvent as a reference. Fluorescence measurements were performed on an Agilent CARY Eclipse fluorescence spectrophotometer. Confocal images were performed on Laser Scanning Confocal Microscope (Andor iQ 3.2) with 100×oil-immersion objective lens (laser combination: 488; 561; 640 nm). Super-resolution images were obtained with Nikon N-SIM 5.0 Super-Resolution Microscope System with a motorized inverted microscope ECLIPSE Ti2-E, a 100×/ NA 1.49 oil immersion TIRF objective lens (CFI HP), and an ORCA-Flash 4.0 SCMOS camera (Hamamatsu Photonics K.K.). Laser combination: 405; 488; 561; 640 nm.

#### *Design and construction of plasmids.*

All plasmid constructions underwent validation through DNA sequencing.

The GLH and CLH constructs were generated using the pET22b-HaloTag7-His6 (synthesized commercially) prokaryotic expression vector as follows: First, sfGFP sequence with 5 amino acids linker added to the C-terminal was PCR-amplified from pET28a-sfGFP (Addgene #85492). mCherry sequence with 4 amino acids linker added to the C-terminal was PCR-amplified from pCMV-mCherry-Tubulin-6 (Addgene #55147). Then, the vector pET22b-HaloTag7-His6 was digested with EcoRI restriction enzyme (TAKARA), the sfGFP or mCherry sequences was connected by One Step Cloning Kit (YEASEN) to obtain pET22b-GLH-His6 and pET22b-CLH-His6 plasmids, respectively.

The CH construct was generated as follows: mCherry sequence deleted 2 amino acids of C-terminal was PCR-amplified from pCMV-mCherry-Tubulin-6 (Addgene #55147). The linearized vector containing the Halo sequence with a deletion of 3 amino acids at the N-terminus was amplified from pET22b-HaloTag7-His6 (Addgene #55147). We finally connected them by One Step Cloning Kit (YEASEN) to obtain pET22b-CH-His6 plasmid.

For transient expression of GLH, CLH, and CH in mammalian cells, we used pCMV-H2B-pSNAPf (NEB #N9186) as the vector. The specific construction was as follows: We first PCR-amplified the GLH, CLH, and CH sequences containing NheI and XhoI restriction sites from pET22b-GLH-His6, pET22b-CLH-His6 and pET22b-CH-His6 plasmids, respectively. Next, the pCMV-H2B-pSNAPf vector was digested with NheI and XhoI restriction enzymes (TAKARA) to remove the SNAP sequence. Finally, the GLH, CLH or CHL sequence was inserted into vector by One Step Cloning Kit (YEASEN) to obtain pCMV-H2B-GLH, pCMV-H2B-CLH, and pCMV-H2B-CH plasmids, respectively.

The construction of CH fused to the mitochondrial target protein was as follows: First, we PCR-amplified the CH sequence containing BamHI and XhoI restriction sites from the pET22b-CH-His6 plasmid. Next, we double-digested the vector pCMV-mApple-TOMM20 (Addgene #54955; TOMM20: mitochondrial outer membrane protein) with BamHI and XhoI restriction enzymes (TAKARA). The CH sequence was then inserted into the C-terminus of the TOMM20 sequence in the vector by One Step Cloning Kit (YEASEN), resulting in the pCMV-TOMM20-CH plasmid.

The construction of CH fused to the endoplasmic reticulum target protein was as follows: First, we PCR-amplified the CH sequence containing NotI and EcoRI restriction sites from the pET22b-CH-His6 plasmid. Next, we double-digested the vector pCDNA3-UnaG-Sec61β (Addgene #83413; Sec61β: endoplasmic reticulum translocon protein) with NotI and EcoRI restriction enzymes (TAKARA). The CH sequence was then inserted into the N-terminus of the Sec61β sequence in the vector by One Step Cloning Kit (YEASEN), resulting in the pCDNA3-CH-Sec61β plasmid.

The construction of CH fused to the lysosome target protein was as follows: First, we PCR-amplified the CH sequence containing NheI and XhoI restriction sites from the pET22b-CH-His6 plasmid. Next, we double-digested the vector pCMV-mCherry-Rab7 (Addgene #55127; Rab7: small GTPase) with NheI and XhoI restriction enzymes (TAKARA). The CH sequence was then inserted into the N-terminus of the Rab7 sequence in the vector by seamless cloning, resulting in the pCMV-Rab7-CH plasmid.

The construction of GLH expressed in mammalian cytoplasm was as follows: First, we PCR-amplified the GLH sequence containing NheI and NotI restriction sites from the pET22b-GLH-His6 plasmid. Next, we double-digested the vector pCMV-mApple-TOMM20 (Addgene #54955) with BamHI and XhoI restriction enzymes (TAKARA). The GLH sequence was then inserted into the vector by the One Step Cloning Kit (YEASEN) to obtain the pCMV-GLH plasmid.

The SLH construct was generated using the pET22b-mStaygold-His6 (synthesized commercially) prokaryotic expression vector as follows: according to the literature, we added the N-terminal region of eGFP (MASTGEELFTGVV, called N1) and the 10 amino acids of the C-terminus of dfGFP (PWHEPSASAV, called C4) to the gene sequence of the pET22b-mStaygold-His6 version by PCR. The introduction of these domains can significantly improve the brightness and stability of mStaygold in cells. In addition, the C-terminal sequence was further amplified to add a 5-amino acid linker to obtain the N1-mStaygold-C4-Linker sequence. At the same time, we amplified the Halo sequence containing NheI and NotI restriction sites from the pET22b-HaloTag7-His6 plasmid. Next, we double-digested the vector pCMV-mApple-TOMM20 (Addgene #54955) with BamHI and XhoI restriction enzymes (TAKARA). The N1-mStaygold-C4-Linker sequence and Halo sequence were then inserted into the vector using a One Step Cloning Kit (YEASEN) to obtain the pCMV-SLH plasmid.

***Protein expression and purification.***

After the plasmid carrying the expression gene was transformed into E. coli strain BL21 (DE3, TransGen Biotech) according to the standard methods. The transformed cells were then plated on selective agar plates containing Ampicillin (Amp). A single colony was inoculated into 5 mL of LB medium containing Amp and incubated overnight at 37°C. The next day, the overnight culture was diluted 1:100 into LB medium (350 mL) with Amp and grown until the optical density ($OD_{600}$) reached 0.6-0.8. Protein expression was induced by adding 0.4 mM isopropyl β-D-1-thiogalactopyranoside (IPTG) and cultured at 18°C for 20 h. After the induction period, the cells were collected by centrifugation and resuspended in lysis buffer (20 mM PBS, 0.3 M NaCl, 1 mM PMSF, pH7.4). The cells were lysed using sonication, and the cell debris was removed by centrifugation to collect the supernatant. The target protein was purified from the lysate using the Ni-NTA resin (GE Healthcare) in 250 mM imidazole buffer (20 mM PBS, 0.3 M NaCl, pH 7.4). Finally, the purified protein was centrifuged using 10 kD ultrafiltration tube (Millipore Corporation) and washed several times with imidazole-free buffer before long-term storage at -80°C. The purity of the protein was analyzed using standard SDS-PAGE. Protein concentration was determined by measuring absorbance at 595 nm (Eppendorf BioPhotometer D30) using Bradford reagent.

***Absorption and fluorescence spectral measurements.***

**O-Rho** and **Si-Rho** were prepared as described in previous work. Stock solutions (1 mM) were prepared in DMSO and diluted to 2 μM in PBS buffer (0.3 M NaCl, pH 7.4) before testing. One aliquot was left untreated, while the other aliquot was incubated at 37°C for 30 minutes after adding the equivalents of HaloTag protein (5 equivalents). Absorption spectra were recorded using an Agilent Cary 60 UV-Vis Spectrophotometer, and fluorescence spectra were recorded using an Agilent Cary Eclipse fluorescence spectrophotometer.

For FRET measurements, GLH and CLH proteins were diluted to 20 μM in PBS buffer (0.3 M NaCl, pH 7.4), labeled with the isotype of **O-Rho** and **Si-Rho** (5 isotypes), and incubated at 37°C for 30 minutes. Excess fluorescent molecules were removed by centrifugation using the 10 kD ultrafiltration tube (Millipore Corporation). Labeled and unlabeled GLH and CLH proteins were diluted to 2 μM in equal amounts, and the fluorescence spectra were measured. The FRET efficiency was calculated using the following formula:

$$E = 1 - \frac{I_{D-A}}{I_D},$$

where $I_D$ is the fluorescence intensity of the donor alone and $I_{D-A}$ is the fluorescence intensity of the donor in the presence of the acceptor.

***Cell culture.***

HeLa (HeLa cyton gartleri, ATCC: CCL-2) cells were purchased from Cell Bank of Type Culture Collection of Chinese Academy of Sciences. HeLa cells were grown in Dulbecco's modified Eagle's medium (DMEM, Gibco) supplemented with 10% fetal bovine serum (FBS, Hyclone) and 1% antibiotics (Hyclone), which were cultured in the humidified atmosphere of 5% $CO_2$ at 37°C. Before imaging, HeLa cells were seeded on glass bottom cell culture dish (Nest, polystyrene, Φ 15 mm) for 24 h to reach 60-80% confluency. The next day, cells were transfected with 0.5 μg of DNA using polyethyleneimine (PEI, Polyscience) according to the manufacturer's protocol. Cell imaging was performed 24-48 hours after transient transfection.

***Confocal fluorescence imaging.***

*FRET fluorescence imaging of genetically encoded fluorophores* (Figure S7, S9a and S10). Cells transiently transfected with pCMV-H2B-GLH, pCMV-H2B-CLH, pCMV-H2B-CH and pCMV-SLH was imaged 24-48 hours after expression. After imaging, the probe (0.5 μM) was added to the cells and incubated for 30 minutes, then washed three times to observe FRET imaging. Laser Scanning Confocal Microscope (Andor iQ 3.2) conditions: 100×oil-immersion objective lens; sfGFP/mStaygold donor channel, λex: 488 nm, collection wavelength: 525±25 nm; **O-Rho** acceptor channel, λex: 561 nm, collection wavelength: 617±73/2 nm; GLH-O/SLH-O FRET channel, λex: 488 nm, collection wavelength: 617±73/2 nm; mCherry donor channel, 561 nm, collection wavelength: 617±73/2 nm; **Si-Rho** acceptor channel, λex: 640 nm, collection wavelength: 683±20 nm; CLH-Si/CH-Si FRET channel, λex: 561 nm, collection wavelength: 683±20 nm. The donor and FRET fluorescence intensity of more than 8-15 cells were analyzed in Andor iQ 3.2 software. The experiment was repeated 3 times.

*No-wash fluorescence imaging of traditional fluorogenic fluorophores* (Figure S2 and S3). Blank Hela cells and Hela cells transiently transfected with pCMV-H2B-Halo were imaged after adding the probe (10-4000 nM) and incubating for 30 minutes. Laser Scanning Confocal Microscope (Andor iQ 3.2) conditions: 100×oil-immersion objective lens; **O-Rho** channel, λex: 561 nm, collection wavelength: 617 ± 73/2 nm; **Si-Rho** channel, λex: 640 nm, collection wavelength: 683 ± 20 nm. The cell experiment was repeated 3 times, and the fluorescence intensity of more than 40-50 cells was analyzed in Andor iQ 3.2 software each time.

*No-wash fluorescence imaging of genetically encoded fluorophores* (Figure 2 and Figure 3). Cells transfected with pCMV-H2B-GLH, pCMV-H2B-CLH, and pCMV-H2B-CH were imaged after adding the probe (50-4000 nM) and incubating for 30 minutes. Laser Scanning Confocal Microscope (Andor iQ 3.2) conditions: 100×oil-immersion objective lens; **O-Rho** channel, λex: 561 nm, collection wavelength: 617±73/2 nm; **Si-Rho** channel, λex: 640 nm, collection wavelength: 683±20 nm; GLH-O channel, λex: 488 nm, collection wavelength: 617 ±73/2 nm; CLH-Si/CH-Si channel, λex: 561 nm, collection wavelength: 683±20 nm. The cell experiment was repeated 3 times, and the fluorescence intensity of 40-50 cells was analyzed in Andor iQ 3.2 software each time.

*Various compartments (nucleus, mitochondria, endoplasmic reticulum, lysosomes) imaging of CH-Si* (Figure S8). Cells transfected with pCMV-H2B-CH, pCMV-Tom20-CH, pCDNA3-CH-Sec61β, pCMV-CH-Rab7 were imaged after adding the **Si-Rho** (0.5 μM) and incubating for 30 minutes. CH-Si channel, λex: 561 nm, collection wavelength: 683±20 nm. The cell experiment was repeated 3 times, and the fluorescence intensity of more than 20 cells was analyzed in Andor iQ 3.2 software each time.

*Dual-color confocal imaging* (Figure S9). HeLa cells co-transfected with pCMV-Tom20-CH and pCMV-mCherry-Rab7 plasmids, then imaged in red and green channel after adding the **Si-Rho** (0.5 μM) and incubating for 30 minutes, respectively.

***Structured illumination microscopy (SIM) imaging.***

*No-wash fluorescence imaging of genetically encoded fluorophores* (Figure 4). Cells transfected with pCMV-H2B-CH were imaged after adding the probe (2000 nM) and incubating for 30 minutes. Nikon N-SIM 5.0 Super-Resolution Microscope conditions: 100×oil-immersion objective lens; **Si-Rho** channel, λex: 640 nm, collection wavelength: 663-738 nm; CH-Si channel, λex: 561 nm, collection wavelength: 700 ± 75/2 nm. The cell experiment was repeated 3 times, and the fluorescence intensity of 40-50 cells was analyzed in software provided with the system each time.

*Four-color imaging* (Figure 4). HeLa cells co-transfected with pCMV-Tom20-CH, pCMV-mCherry-Rab7 and pCMV-Sec61β-EGFP plasmids were incubated with **Si-Rho** (0.5 μM), and Hoechst 33342 (0.5 μM) for 60 minutes, and then imaged in pink, orange, green and blue channels respectively. Super-Resolution Microscope conditions: 100×oil-immersion objective lens; blue channel, λex: 405 nm, collection wavelength: 435-485 nm; green channel, λex: 488 nm, collection wavelength: 500-545 nm; orange channel, λex: 561 nm, collection wavelength: 570-640 nm; pink channel, λex: 561 nm, collection wavelength: 700 ± 75/2 nm.

*Mitochondrial dynamic imaging* (Figure 5). Mitochondrial and endoplasmic reticulum interaction imaging: HeLa cells co-transfected with pCMV-Tom20-CH, and pCMV-EGFP-Sec61β plasmids, and added the **Si-Rho** (0.5 μM) and incubating for 30 minutes. Then, the cells were co-imaged using the red and green channels every 10 seconds for continuously 20 min. Super-Resolution Microscope conditions: 100×oil-immersion objective lens; green channel, λex: 488 nm, collection wavelength: 500-545 nm; pink channel, λex: 561 nm, collection wavelength: 700 ± 75/2 nm.

Mitochondrial and lysosome interaction imaging: HeLa cells co-transfected with pCMV-Tom20-CH and pCMV-mCherry-Rab7 plasmids, and added the **Si-Rho** (0.5 μM) and incubating for 30 minutes. Then, the cells were co-imaged using the red and green channels every 10 seconds for continuously 20 min. Super-Resolution Microscope conditions: 100×oil-immersion objective lens; green channel, λex: 561 nm, collection wavelength: 570-640 nm; pink channel, λex: 561 nm, collection wavelength: 700 ± 75/2 nm.

*Photobleaching kinetics imaging* (Figure 6). For SIM photobleaching images of GLH/SLH proteins, imaging was conducted with a 488 nm laser at 30% power (0.16 kW/cm$^2$, collection wavelength: 605 ± 35 nm) and 200 ms exposure time. A total of 121 frames were recorded within 20 minutes with a 10-second interval between each frame. The final SIM images were displayed with the same intensity levels.

***Data processing and analysis.***

The generation of all spectral curves in this study was processed using Origin 2018 software. Fluorescence intensity data of live cell imaging were obtained by the built-in analysis software of the imaging system. Signal-to-noise ratio was calculated using GraphPad Prism 8 and Origin2018 software. Live cell images were analyzed in ImageJ Fiji.

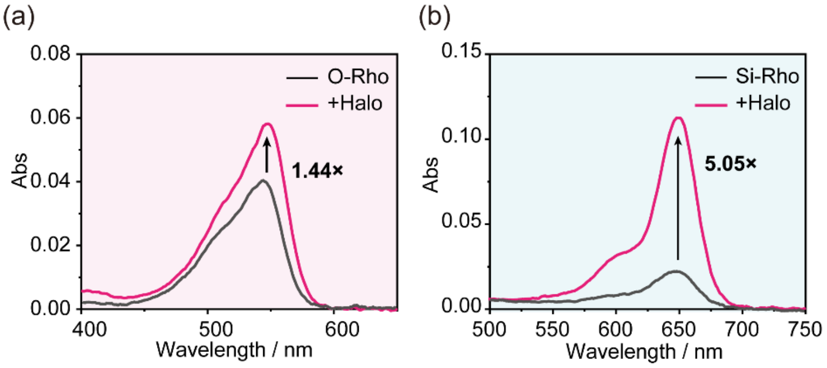

**Figure S1.** (a, b) Absorption spectra of **O-Rho** and **Si-Rho** (2 μM) in PBS buffer (pH 7.4) measured in the presence (pink line) and absence (black line) of HaloTag (10 μM). The numbers indicate the ratio of **O-Rho** absorption at 544 nm and **Si-Rho** absorption at 648 nm.

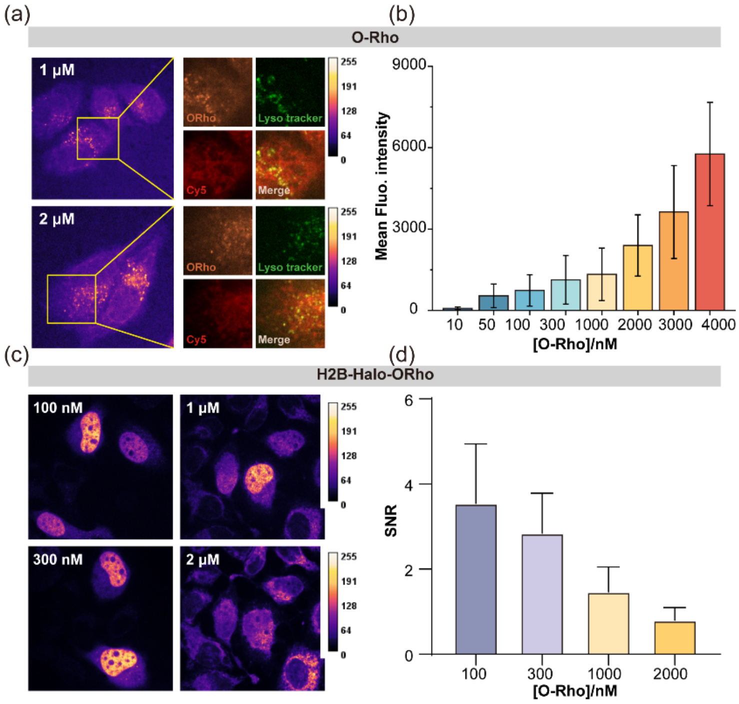

**Figure S2.** (a) No-wash imaging of blank Hela cells labeled with **O-Rho** (1-2 μM). The right panel of images shows the co-localization imaging of **O-Rho** (orange) with mitochondria (stained with Cy5, red) and lysosomes (stained with Lyso-tracker green, green). (b) Statistical analysis of fluorescence intensity of blank HeLa cells stained with **O-Rho**. The graph displays the average ± SD from more than 50 cell experiments. (c) No-wash imaging of H2B-Halo protein labeled with varying concentrations **O-Rho** (0.1-2 μM). Calibration bar shows the normalized fluorescence intensity. (d) Statistical analysis of the SNR in HeLa cells expressing

H2B-Halo proteins stained with **O-Rho** ($\lambda_{ex}$: 561 nm, collected: 617±73/2 nm). The graph displays the average ± SD from more than 50 cell experiments.

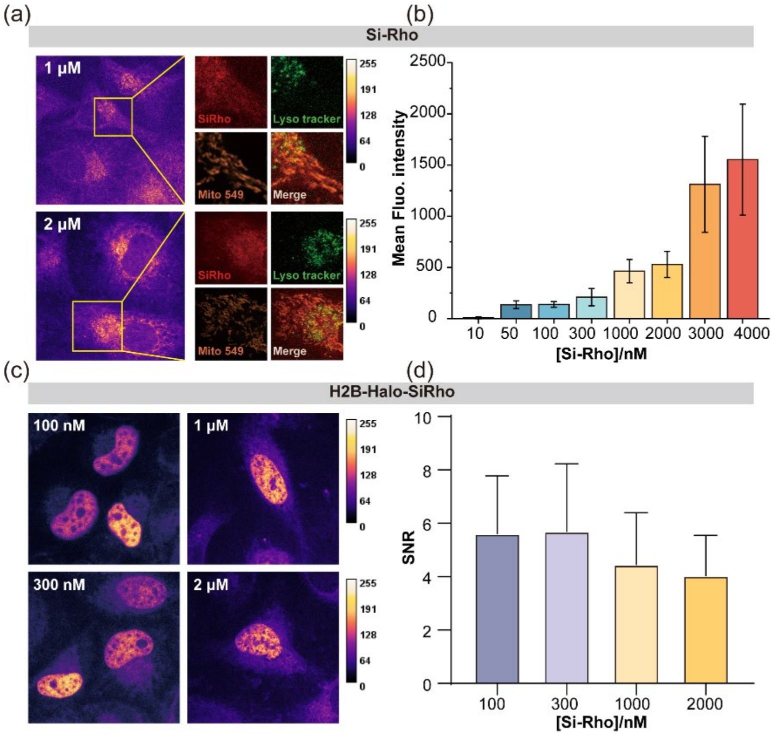

**Figure S3.** (a) No-wash imaging of blank Hela cells labeled with **Si-Rho** (1-2 μM). The right panel of images shows the co-localization imaging of **Si-Rho** (orange) with mitochondria (stained with Cy5, red) and lysosomes (stained with Lyso-tracker green, green). (b) Statistical analysis of fluorescence intensity of blank HeLa cells stained with **Si-Rho**. The graph displays the average ± SD from more than 50 cell experiments. (c) No-wash imaging of H2B-Halo protein labeled with varying concentrations **Si-Rho** (0.1-2 μM). Calibration bar shows the normalized fluorescence intensity. (d) Statistical analysis of the SNR in HeLa cells expressing H2B-Halo proteins stained with **Si-Rho** ($\lambda_{ex}$: 640 nm, collected: 683±20 nm). The graph displays the average ± SD from more than 50 cell experiments.

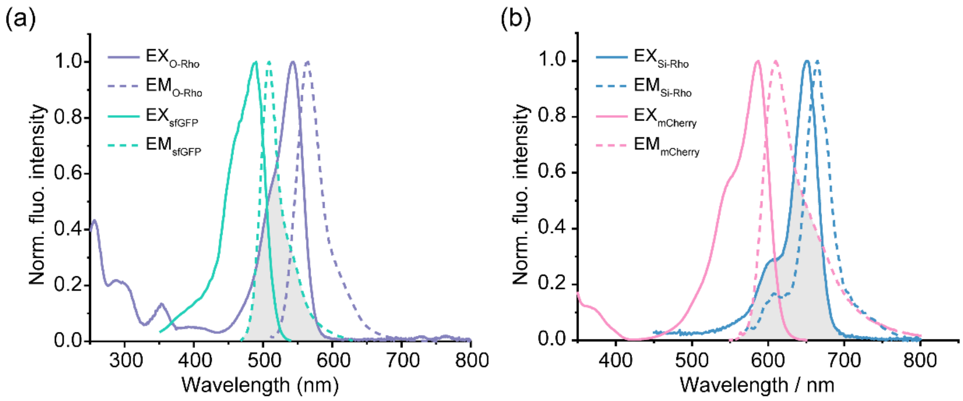

(c)

| | $\lambda_{ex}$ nm | $\lambda_{em}$ nm | ε $M^{-1}\cdot cm^{-1}$ | Φ | Turn on $A/A_0$ | Turn on $F/F_0$ |
|---|---|---|---|---|---|---|
| O-Rho | 540 | 565 | 63680 | 0.85 | 1.44 | 1.6 |
| sfGFP | 488 | 510 | 83300 | 0.65 | - | - |
| Si-Rho | 643 | 662 | 141000 | 0.41 | 5.05 | 6.2 |
| mCherry | 587 | 610 | 72000 | 0.22 | - | - |

**Figure S4.** (a, b) Normalized excitation (EX) and emission (EM) spectra of donor sfGFP (a) and mCherry (b), along with acceptor dye **O-Rho** and **Si-Rho**. The grey region indicates the overlap between the emission spectrum of the donor and the excitation spectrum of the acceptor. (c) Comparison of the properties between fluorescent proteins and Rho dyes.

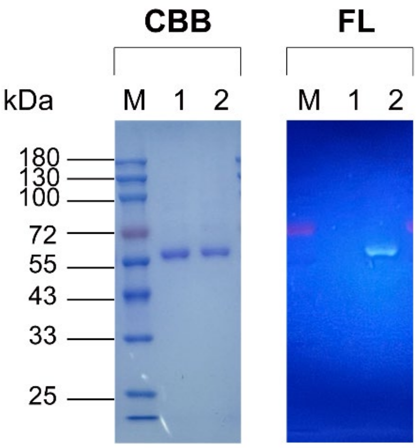

**Figure S5.** The SDS-PAGE analysis of GLH protein labeled and non-labeled with NAP488 dye. The gel was stained with Coomassie Brilliant Blue (CBB) for total protein visualization and scanned for fluorescence (FL) under UV excitation. Lanes displayed from left to right as follows: (M) Prestained Protein Marker ranging from 8-180 kDa, (1) GLH, and (2) GLH-NAP488.

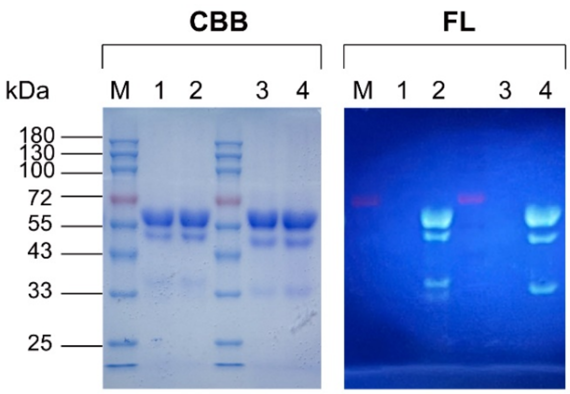

**Figure S6.** The SDS-PAGE analysis of CLH and CH proteins labeled and non-labeled with NAP488 dye. The gel was stained with Coomassie Brilliant Blue (CBB) for total protein visualization and scanned for fluorescence (FL) under UV excitation. Lanes displayed from left to right as follows: (M) Prestained Protein Marker ranging from 8-180 kDa, (1) CLH, (2) CLH-NAP488, (3) CH, and (4) CH-NAP488. Smaller bands observed in all lanes are attributed to hydrolysis of RFP-like chromophores under denaturing conditions of SDS-PAGE.

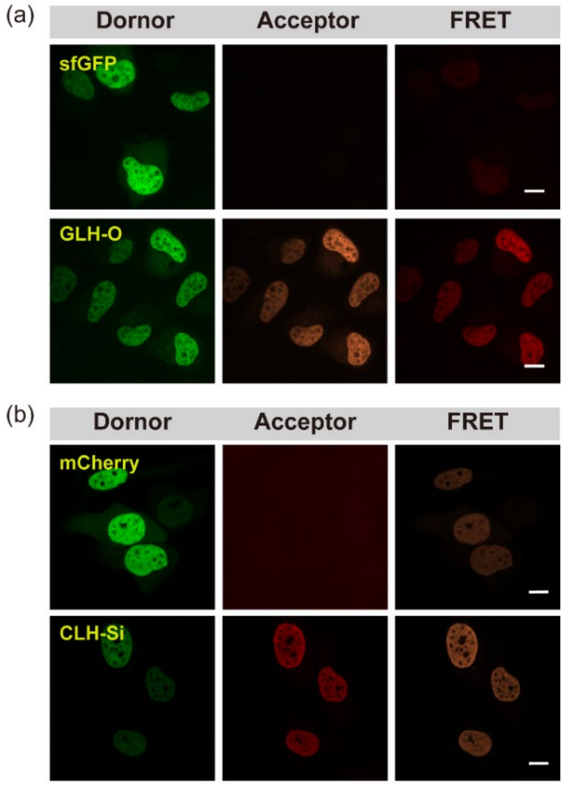

**Figure S7.** (a-b) Confocal fluorescence images of HeLa cells expressing H2B-GLH (a) and H2B-CLH (e) labeled and unlabeled with **O-Rho** or **Si-Rho**, were captured in the donor channel, acceptor channel and FRET channel, respectively. The images in each channel were adjusted with the same contrast. Scale bar: 10 µm.

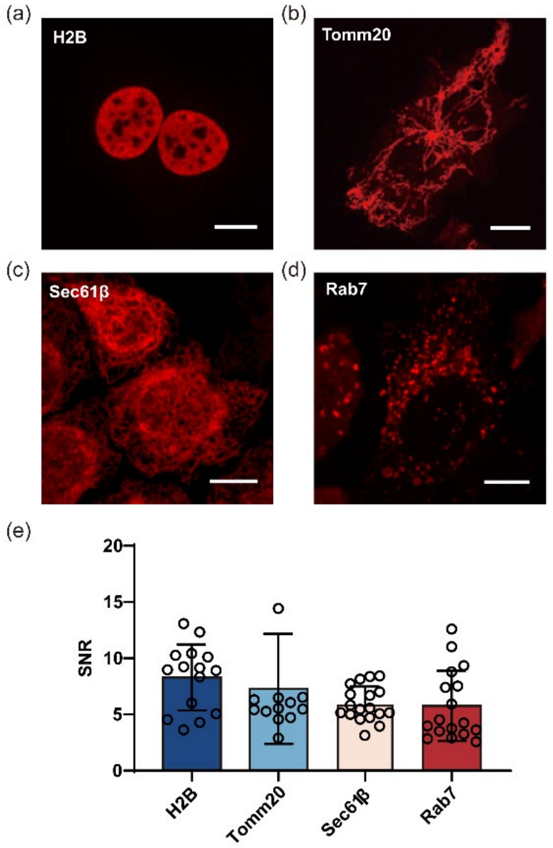

**Figure S8.** Confocal fluorescence images of the nucleus (a), mitochondria (b), endoplasmic reticulum (c), and lysosome (d) organelles in Hela cells using CH-Si probe. Excitation: 561 nm. Emission: 663-703 nm. Scale bar: 10 µm. (e) SNR analysis of CH-Si in imaging of different cell organelles. The graph displays the average values ± SD from 20-30 cell experiments.

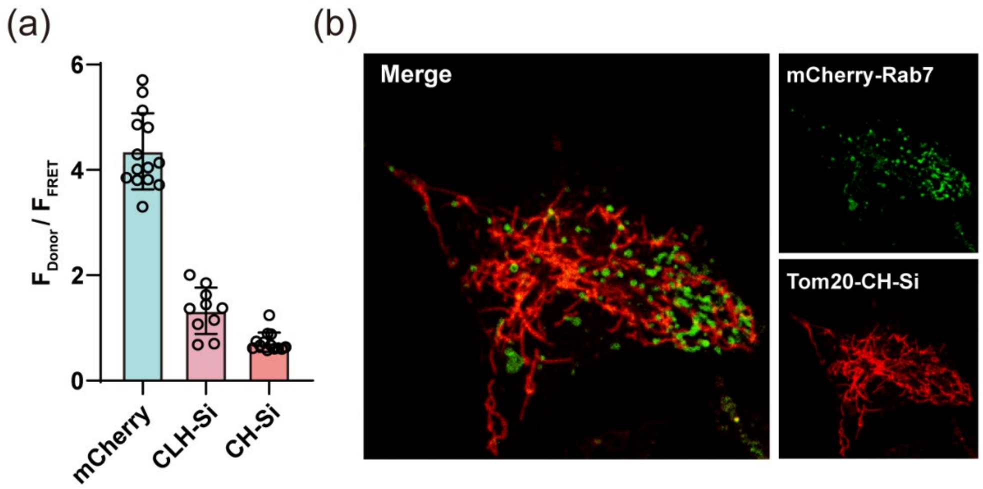

**Figure S9.** (a) Fluorescence intensity ratio in the donor channel and FRET channel from confocal fluorescence imaging of mCherry, CLH-Si and CH-Si. The graph shows the mean ± SD from 20-30 cells in three independent experiments. (b) Two-color confocal image of Rab7 labeled with mCherry (green) and Tom20 labeled with CH-Si (red).

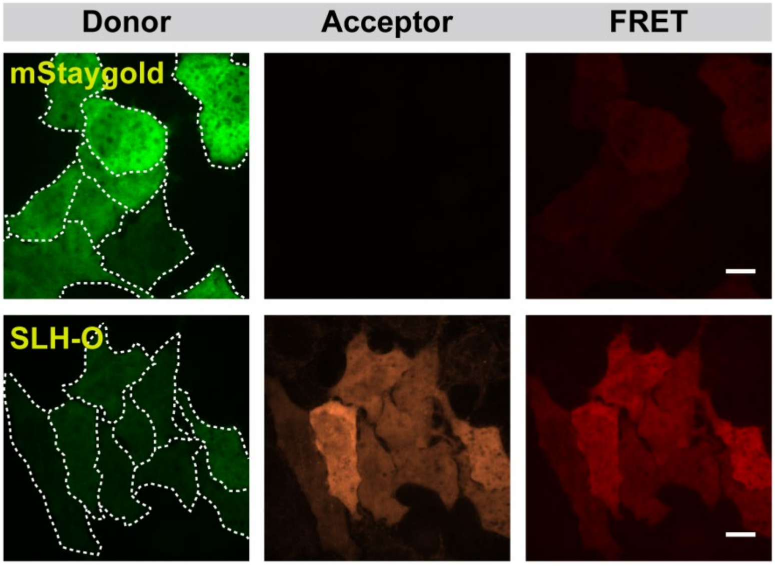

**Figure S10.** Confocal fluorescence images of HeLa cells expressing mStaygold and SLH labeled and unlabeled with **O-Rho**, were captured in the donor channel, acceptor channel and FRET channel, respectively. The images in each channel were adjusted with the same contrast. Scale bar: 10 μm.

**Table S1.** Amino acid sequences and molecular weights of the linker regions in GLH, SLH, CLH, and CH proteins.

| | sfGFP-Linker-Halo Protein squence | MW (Da) |
|---|---|---|
| GLH | ……ITHGMDELYKGSAFKMAEIGTGFPFDPHYVE…… | 62000 |
| | mStaygold-Linker-Halo Protein squence | MW (Da) |
| SLH | ……WHEPSASAVGSAFK MAEIGTGFPFDPHYVE…… | 60420 |
| | mCherry-Linker-Halo Protein squence | MW (Da) |
| CLH | ……STGGMDELYKSGLRMAEIGTGFPFDPHYVE…… | 62860 |
| CH | ……STGGMDELIGTGFPFDPHYVE…… | 61830 |
